# Behavioral and brain signatures of substance use vulnerability in childhood

**DOI:** 10.1101/2020.10.19.346403

**Authors:** Kristina M. Rapuano, Monica D. Rosenberg, Maria T. Maza, Nicholas Dennis, Mila Dorji, Abigail S. Greene, Corey Horien, Dustin Scheinost, R. Todd Constable, BJ Casey

## Abstract

The prevalence of risky behavior such as substance use increases during adolescence; however, the neurobiological precursors to adolescent substance use remain unclear. Predictive modeling may complement previous work observing associations with known risk factors or substance use outcomes by developing generalizable models that predict early susceptibility. The aims of the current study were to identify and characterize behavioral and brain models of vulnerability to future substance use. Principal components analysis (PCA) of behavioral risk factors were used together with connectome-based predictive modeling (CPM) during rest and task-based functional imaging to generate predictive models in a large cohort of nine- and ten-year-olds enrolled in the Adolescent Brain & Cognitive Development (ABCD) study (NDA release 2.0.1). Dimensionality reduction (*n*=9,437) of behavioral measures associated with substance use identified two latent dimensions that explained the largest amount of variance: risk-seeking (PC1; e.g., curiosity to try substances) and familial factors (PC2; e.g., family history of substance use disorder). Using cross-validated regularized regression in a subset of data (Year 1 Fast Track data; n>1,500), functional connectivity during rest and task conditions (resting-state; monetary incentive delay task; stop signal task; emotional n-back task) significantly predicted individual differences in risk-seeking (PC1) in held-out participants (partial correlations between predicted and observed scores controlling for motion and number of frames [r_p_]: 0.07-0.21). By contrast, functional connectivity was a weak predictor of familial risk factors associated with substance use (PC2) (r_p_: 0.03-0.06). These results demonstrate a novel approach to understanding substance use vulnerability, which—together with mechanistic perspectives—may inform strategies aimed at early identification of risk for addiction.

## Introduction

Although adolescents are prone to a variety of risky behaviors, experimentation and use of substances are particularly common amongst youth—with nearly a quarter of 8th graders and half of 12th graders reporting experimentation with illicit substances (Johnston et al., 2018). The consequences of substance use range from accident and injury to the individual to economic strain, with the overall cost associated with alcohol, tobacco, and illicit substances in the US estimated to exceed $740 billion annually (NIDA, 2017). An extensive body of work has characterized the impact of substances on brain function in individuals that have already begun to use or are dependent on substances (Balodis & Potenza, 2015; Garavan et al., 2000; Houck et al., 2013; Janes et al., 2012; Ma et al., 2010, 2011; Tapert et al., 2004); however, less is known about the impact of potential risk factors for substance use on the developing brain and how these precursors may motivate susceptible individuals to begin using substances early in life.

Considerable variability in the propensity to engage in risky behaviors has been observed across adolescents in the real world (Johnston et al., 2018) and in the lab (Galvan et al., 2007). These individual differences in behavior—such as substance use—may be explained in part by the impact of environmental and genetic risk factors on developing brain circuitry. For example, early experimentation with substances in youth has consistently been shown to correlate with later outcomes. Early sips of alcohol (before 6th grade) have been linked to earlier onset of drinking a full glass of alcohol (Donovan & Molina, 2011), being drunk by 9th grade (Donovan & Molina, 2011; Jackson et al., 2015), alcohol misuse by ages 17-18 (Hawkins et al., 1997), and alcohol dependence later in life (Grant & Dawson, 1997; Prescott & Kendler, 1999). Similarly, an individual’s age at first cigarette puff and first full cigarette has been associated with a higher likelihood of later identification as a smoker (Azagba et al., 2015). Finally, early thoughts about intention and curiosity to use substances have been shown to relate to experimentation and later alcohol and tobacco use in youth (Andrews et al., 2003; Pierce et al., 1996; Trinidad et al., 2017), and adolescents that experiment with one substance early on tend to concurrently experiment or engage with others (e.g., marijuana and tobacco; (Okoli et al., 2008).

In addition to curiosity and early experimentation, several familial factors have been shown to increase risk for later substance use in children. For example, family history of alcohol dependence is associated with problem drinking in college students (Labrie et al., 2010)—independent of age of first drink (Grant, 1998; Prescott & Kendler, 1999)—with the heritability of alcoholism estimated to be between 50-60% (McGue, 1999). Similarly, family history of drug use has been associated with drug use severity (Pickens et al., 2001), with an estimated heritability of drug dependence ranging from 0.39 (for hallucinogens) to 0.72 (for cocaine) (Goldman et al., 2005), and children with parents who smoke at home are more likely to begin or desire to begin smoking early in life (Gilman et al., 2009; C. Jackson & Henriksen, 1997). Despite these well-established behavioral associations, the extent to which vulnerability is represented in the developing brain prior to observable phenotypic differences in behavior remains unclear.

Previous work has highlighted differences in brain circuits implicated in reward motivation and cognitive control in individuals with addiction or at risk for addiction (Balodis & Potenza, 2015; Bjork et al., 2004; Whelan et al., 2012; Yau et al., 2012). For example, neuroimaging studies have observed differences during a monetary reward task in adults with addiction (Balodis & Potenza, 2015) as well as in those genetically at-risk for addiction (Villafuerte et al., 2012; Yau et al., 2012). Moreover, brain responses during impulse control tasks have been consistently associated with substance use in adolescence (Mahmood et al., 2013; Norman et al., 2011; Tapert et al., 2007; Wetherill et al., 2013; Whelan et al., 2012). The development of these circuits has been suggested to be hierarchical, such that brain circuitry governing reward motivation shows functional and structural changes prior to changes in cognitive control circuitry (Casey et al., 2019; Heller et al., 2016). This differential development of reward- and control-related circuitry is considered to encourage exploratory behavior and allow adolescents to adapt to new environmental challenges (Casey, 2015), and also plays a role in heightened sensation seeking (Casey & Jones, 2010; Steinberg, 2008). Thus, children and adolescents may be particularly susceptible to substance-related cues and risk factors promoting early initiation.

Beyond heightened vulnerability attributed to normative neurodevelopment, differences in a variety of risk factors (e.g., psychopathology, behavioral tendencies) may impact the developing brain in such a way that further promotes substance use. A growing literature has begun to characterize vulnerability to substance use by examining regional brain differences between individuals considered to be at risk relative to those at a lower risk. Due to the high heritability of substance dependence, “risk” has commonly been defined by genetic risk or family history of substance use disorder. For example, adolescents with a family history of alcohol use disorder exhibit differences in striatal activity during tasks associated with response inhibition (Heitzeg et al., 2010) and monetary reward (Yau et al., 2012), as well as differential prefrontal activity during tasks associated with reward (Cservenka & Nagel, 2012; Ivanov et al., 2012), working memory (Cservenka et al., 2012), and inhibition (Acheson et al., 2014). Other studies have identified neurobiological markers of vulnerability by examining prospective relationships with known outcomes. Using longitudinal approaches, adolescent substance use initiation and severity has been predicted by heightened striatal activity during monetary reward tasks (Morales et al., 2018; Stice et al., 2013), reduced activation in regions associated with response inhibition (Norman et al., 2011), and reduced frontoparietal activity during a working memory task (Squeglia et al., 2012; Tervo-Clemmens et al., 2018). Recent work has sought to reconcile findings in the literature based on disparate ways of defining and investigating vulnerability for substance use. In a metaanalysis, (Tervo-Clemmens et al., 2020) observed reliable differences associated with substance use vulnerability in the striatum, and this activation pattern was consistent between groups of individuals that were identified based on family history and those that were prospectively identified as at-risk, suggesting the involvement of common underlying circuitry.

Given the growing interest in identifying biomarkers of substance use vulnerability (Heitzeg et al., 2015; Squeglia & Cservenka, 2017), predictive models that precede initiation as well as knowledge of future outcomes will be an important gap to address. Although studies characterizing vulnerability in the brain provide important insight into the mechanisms underlying substance use, these primarily correlational findings are limited in terms of their generalizability and out-of-sample prediction. There has been a recent push towards emphasizing predictive approaches as a way of understanding behavior—in addition to more commonly practiced descriptive-based approaches (Gabrieli et al., 2015; Rosenberg et al., 2018; Varoquaux & Poldrack, 2019; Woo et al., 2017; Yarkoni & Westfall, 2017). Rather than explaining regional brain differences attributable to specific risk factors or task conditions, a predictive model of substance use would capture a more nuanced and generalizable representation of behavioral and neurobiological vulnerability. Predictive models of vulnerability, constrained by hypothesis-driven findings derived from mechanistic approaches, may have a greater potential to benefit early intervention and prevention strategies.

Connectome-based predictive modeling (CPM) has proven to be fruitful in generating predictions about behavior both across individuals (Finn et al., 2015; Rosenberg et al., 2016a; Shen et al., 2017) and within individuals over time (Lichenstein et al., 2019; Rosenberg et al., 2020; Yip et al., 2019a), and has specifically been suggested to provide biomarkers for predicting addiction (Yip et al., 2019b). Because of the circuit-based changes that occur during development, CPM may be especially informative for characterizing brain predictors of vulnerability to future substance use in children. Functional connectivity at rest not only distinguishes an individual (i.e., represents a neural “fingerprint”) (Finn et al., 2015) but also reflects developmental changes in brain circuitry (Dosenbach et al., 2010; Nielsen et al., 2019) as well as genetic and environmental history (Ge et al., 2017; Miranda-Dominguez et al., 2018; Thompson et al., 2013), and has been highlighted as an effective measure of circuit-based imbalances in addiction (Fedota & Stein, 2015; Sutherland et al., 2012). Moreover, functional connectivity during cognitive challenges has been shown to improve behavioral predictions when compared to functional connectivity during rest (Greene et al., 2018; Rosenberg et al., 2016a; Yoo et al., 2018). This distinction may point towards the usefulness of in-scanner tasks for assessing circuitry associated with substance use, such as circuits involved in reward and inhibitory control processing (Balodis & Potenza, 2015; Bjork et al., 2004; Whelan et al., 2012; Yau et al., 2012) as well as general cognitive function (Squeglia et al., 2012). Thus, functional brain connectivity during rest and during task are strong candidate measures for predicting vulnerability to future use in the developing brain prior to onset of use.

Here we use a large-scale open-access dataset, the Adolescent Brain and Cognitive Development (ABCD) study (Casey et al., 2018), to characterize brain and behavioral signatures of vulnerability for future substance use during childhood. As one of the primary objectives of the ABCD study is to understand the impact of substance use on the developing brain and neurocognitive outcomes (https://abcdstudy.org), this study incorporates a comprehensive battery of substance-related questionnaires and surveys (Lisdahl et al., 2018). The current analysis utilizes this resource to 1) quantify behavioral dimensions of vulnerability for future substance use based on this battery; and 2) identify brain signatures that predict individual differences among these dimensions during a range of cognitive conditions. Given previous literature exploring behavioral and neurobiological vulnerability to substance use, as well as previous work using connectome-based prediction, we hypothesize that functional connectivity during rest and task will predict individual differences in latent dimensions of substance use vulnerability. Using a combination of data-driven and hypothesis-driven approaches, we show that functional connectivity in the developing brain predicts vulnerability to substance use in children prior to initiation.

## Methods

### Participants

Participants in the current study are enrolled in the ongoing Adolescent Brain Cognitive Development (ABCD) Study—a 10-year longitudinal study that aims to further our understanding of the environmental and genetic influences on brain development and their roles in substance use and other health outcomes (Garavan et al., 2018). This large-scale study tracks 11,875 children between the ages of 9-11 recruited from 21 research sites across the United States. Although designed to approximate the diversity of the US population based on sex, race, ethnicity, and socioeconomic status (Compton et al., 2019), the population characteristics of the sample described in the current analyses are not guaranteed to match national estimates (Garavan et al., 2018).

Parental informed consent and child assent were obtained from all participants and approved by centralized and institutional review boards at each data collection site. Study-wide exclusionary criteria included a diagnosis of schizophrenia, moderate to severe autism spectrum disorders, intellectual disabilities, or substance use disorders at the time of recruitment. Additionally, children with major and persistent neurological disorders, multiple sclerosis, sickle cell disease or certain seizure disorders such as Lennox-Gastaut syndrome, Dravet syndrome, and Landau Kleffner syndrome were excluded.

The current study includes year-one (baseline) assessments from the curated 2.0.1 release of the ABCD study data set. Analyses utilize behavioral data collected on the full baseline cohort (*n*=11,875) and all fMRI data available through ABCD’s Fast Track option as of April 2018 (*n*=5,772). Children with a mild autism spectrum diagnosis or any history of epilepsy were excluded from the current analysis. Low quality and/or high-motion fMRI data (described in *Methods: preprocessing*) were additionally excluded from neuroimaging analyses. To avoid confounds associated with family relatedness, behavioral and neuroimaging analyses were run with and without the inclusion of siblings.

### Study Design

#### Psychological and behavioral measures

Parent- and child-reported psychological and behavioral assessments associated with substance use were compiled for inclusion in the present study. To constrain the current analysis to variables directly associated with substances, behavioral measures were selected by querying the full battery of ABCD baseline measures for items explicitly referring to substances (e.g., “alcohol”, “substances”; Table S1). Itemized scores were collapsed across substances where applicable in order to identify risk for general substance use rather than for specific substances.

#### Resting state fMRI data collection

ABCD images were acquired using Siemens Prisma, Philips, or GE 750 3T scanners with a 32-channel head coil. Detailed acquisition parameters have been previously described in the literature (Casey et al., 2018). Scan sessions included a high-resolution anatomical scan, diffusion weighted images, T2-weighted spin echo images, resting-state fMRI, and task-based fMRI. Functional images were collected through 60 slices in the axial plane using echo-planar imaging sequence with the following parameters: TR = 800 ms, TE = 30 ms, flip angle = 52°, voxel size = 2.4 mm^3^, multiband slice acceleration factor = 6.

Participants completed up to four runs of 5-minute resting-state fMRI scans. ABCD sites with Siemens scanners used Framewise Integrated Real-time MRI Monitoring (FIRMM; (Dosenbach et al., 2017), which monitors head motion in real-time and allows for the discontinuation of resting-state data collection after three runs if 12.5 minutes of low-motion data had been collected.

#### Monetary incentive delay

The monetary incentive delay task (MID) measures components of reward processing including anticipation and outcome of rewards and losses as well as the motivation to gain rewards and mitigate losses (Knutson et al., 2000). MID data were collected in two 50-trial fMRI runs each lasting approximately 5.5 minutes (403 volumes per run after discarded acquisitions). In each trial, participants were presented with an incentive cue (1500-4000 ms) indicating whether they could win $0.20 or $5.00, lose $0.20 or $5.00, or if no money was at stake (Figure 1B). This incentive cue was followed by a fixation delay and then a dynamically-manipulated target lasting 150-500 ms, customized to ensure participants achieved approximately 60% accuracy. In order to achieve the outcome, participants were required to respond during the target presentation. Feedback of reward or loss was provided for each trial.

**Figure 1.**
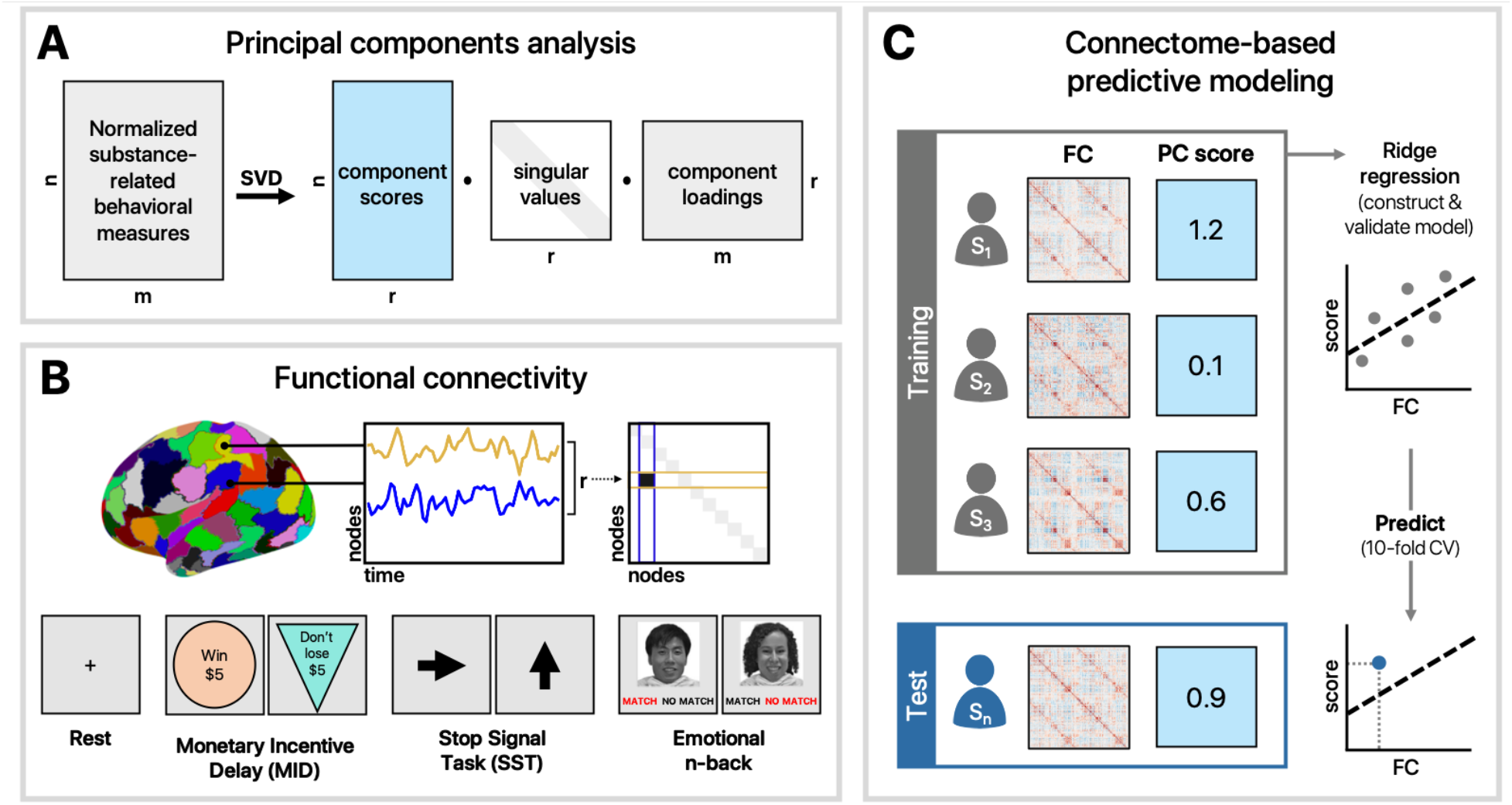
Analysis schematic. **A)** Behavioral measures associated with substance use were normalized and reduced using principal components analysis. **B)** Functional connectivity matrices were generated for each task and rest using a 268-node brain atlas (X. Shen et al., 2013). **C)** A 10-fold cross-validation procedure was used to train and test a ridge regression model to predict individual differences in PCA scores.

#### Stop Signal Task

The Stop Signal Task measures impulsivity and impulse control (Logan, 1994). SST data were collected in two 180-trial fMRI runs each lasting approximately 6-minutes (437 volumes after discarded acquisitions). In each trial, participants were presented with a go signal, represented by a left or right arrow, which they responded to by indicating the direction in which the arrow was pointing with a left or right button press. A stop signal, indicated by an upright arrow, followed 16.67% of the go signals such that participants were required to inhibit their already-initiated response. The timing between go and stop signals was dynamically-manipulated for each subject to ensure approximately 50% accuracy.

#### Emotional n-Back Task

The emotional n-back Task (Cohen et al., 2016a; Cohen et al., 2016b) measures working memory by manipulating cognitive load and emotional processes by using emotive faces in addition to places as stimuli (Barch et al., 2013; Casey et al., 2018). Emotional n-back data were collected in two 80-trial fMRI runs each lasting approximately 5-minutes (403 volumes after discarded acquisitions). Each run consisted of eight task blocks (four 0-back; four 2-back) and four fixation blocks (15 s each). For the 0-back condition, participants were instructed to respond “match” when the current picture was the same as the target picture shown at the start of the block, “no match” if not. For the 2-back condition, participants were instructed to respond “match” when the current picture was the same as the one shown two pictures back, “no match” if not. In each task block, participants were first presented with an instruction cue (2500 ms) to indicate which task condition would follow—0-back (“Target=“ and a picture of the target stimulus) or 2-back (“2-back”). To signal a switch between task conditions, a colored fixation cross (500 ms) preceded each block’s instruction cue. The instruction cue was followed by the interstimulus interval fixation (500 ms) and then a stimulus trial (2500 ms)—a picture of a face (happy, fearful, or neutral expressions) or a place immediately followed by a fixation cross. Each task block consisted of ten trials, 2 of which were targets, 2-3 were non-target lures (repeated incorrect stimuli), and the remainder were non-lures (non-repeated stimuli).

### Behavioral analyses

Data from 287 participants with epilepsy or autism diagnoses were excluded from the 11,431 participants with complete substance use data, resulting in 11,144 participants. Of these, 9,437 remained after the exclusion of siblings (Table 1; Figure S2).

**Table 1.**
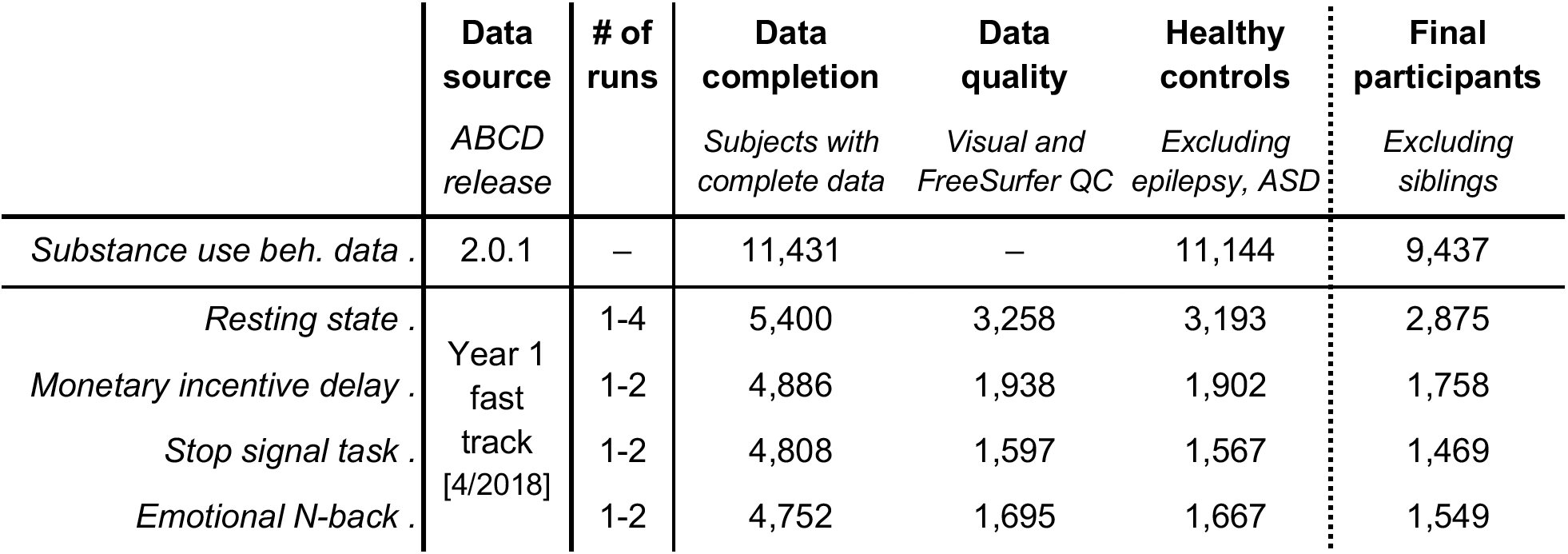
Participant selection. Full baseline data from ABCD Release 2.0.1 were used for behavioral analyses. Year 1 fast track data (available as of April 2018) were used for fMRI analyses. Data were downloaded approximately halfway through baseline data collection. The total number of subjects meeting inclusion criteria for each analysis varied across conditions. All analyses were performed with and without the inclusion of siblings to account for family relatedness.

|  | Data source<br>*ABCD release* | # of runs | Data completion<br>*Subjects with complete data* | Data quality<br>*Visual and FreeSurfer QC* | Healthy controls<br>*Excluding epilepsy, ASD* | Final participants<br>*Excluding siblings* |
| --- | --- | --- | --- | --- | --- | --- |
| *Substance use beh. data* | 2.0.1 | – | 11,431 | – | 11,144 | 9,437 |
| *Resting state* | Year 1 fast track [4/2018] | 1-4 | 5,400 | 3,258 | 3,193 | 2,875 |
| *Monetary incentive delay* |  | 1-2 | 4,886 | 1,938 | 1,902 | 1,758 |
| *Stop signal task* |  | 1-2 | 4,808 | 1,597 | 1,567 | 1,469 |
| *Emotional N-back* |  | 1-2 | 4,752 | 1,695 | 1,667 | 1,549 |

Associations among raw measures related to substance use were first examined using Pearson correlation and visualized using multidimensional scaling (MDS). To better understand the variability in the data, principal components analysis (PCA) was used to reduce the dimensionality of the data (Figure 1A). Because of the skewness and non-normality of the behavioral data (i.e., fewer participants endorsing questions related to substance use as severity increases), summarized data across all substance-related measures were transformed and standardized using the Yeo-Johnson normalizing transformation in *scikit learn* (Pedregosa et al., 2011), which uses maximum likelihood estimates to stabilize variance and minimize the skewness of the data (Yeo & Johnson, 2000). The dimensionality of the normalized data was reduced using singular value decomposition via *scikit learn’s* PCA function. The number of components was determined using Bayesian model selection (Minka, 2001), yielding 14 principal components. The reliability of component loadings was assessed by computing 95% confidence intervals across 10,000 bootstraps. Loadings whose confidence intervals did not cross zero were considered reliable. Normalized substance use data were projected onto the resulting space to produce component scores for each participant.

### Neuroimaging

#### Preprocessing

Raw dicom images for 5,772 participants were downloaded via ABCD Fast Track (April 2018) and preprocessed using BioImage Suite (Joshi et al., 2011) using an approach described in detail elsewhere (Greene et al., 2018; Horien et al., 2019). T1-weighted anatomical images were skull stripped using optiBET (Lutkenhoff et al., 2014)—a modified version of FSL’s brain extraction tool (Smith, 2002), and non-linearly registered to MNI stereotaxic space using B-spline free form deformation. Resulting images were visually inspected to verify the quality of previous preprocessing steps and to ensure that the images were artifact-free. Participants that did not pass visual inspection (*n*=48) were excluded from further analysis, as well as scanning runs with more than 0.15-mm mean frame-to-frame displacement, 2-mm maximum displacement, or more than 3-degrees of rotation. Functional images were realigned to correct for motion, registered to MNI space, and anatomically parcellated using a 268-node whole-brain atlas (Shen et al., 2013). Covariates of no interest were regressed from the data, including linear, quadratic, and cubic drifts, 24-motion parameters (Satterthwaite et al., 2013), mean cerebral-spinal fluid signal, mean white matter signal, and overall global signal. Data were temporally smoothed with a Gaussian filter, σ = 1.95 (approximate cut-off frequency of 0.12 Hz). Pearson correlation coefficients between time courses for every pair of nodes were computed and Fisher *z*-transformed, resulting in a 268×268 functional connectivity matrix for each run and each participant (Figure 1B). After additionally excluding those with low quality anatomical images by FreeSurfer (ABCD NDA name: *fsqc_qc*), data for 3,193 participants remained in the resting-state condition; 1,902 participants for the MID task; 1,567 participants for the SST; and 1,667 participants for the emotional n-back task (Table 1; Figure S3).

#### Predictive modeling

Rest- and task-based functional connectivity matrices from participants meeting inclusion criteria were used to generate predictive models of substance risk-seeking (PC1) and familial risk for substance use (PC2) (Figure 1C). For a given scan condition, connectivity matrices were vectorized and entered into a linear regression model with L2 regularization (i.e., ridge regression) used to predict component scores in the held-out data using 10-fold cross-validation. Component loadings were recomputed in a subset of participants independent from those contributing to a given neuroimaging analysis, such that resulting PCs were consistent with those observed in the behavioral analysis (r > 0.9) but statistically independent from the neuroimaging sample. Resulting loadings were subsequently used to transform substance use data in participants used for connectome-based model generation. To avoid biasing the test set, edge strengths were z-scored across subjects within the training set (i.e., 90% of participants) and the corresponding transformation was subsequently applied to the test set (i.e., 10% of participants) on each fold.

Regression models were developed and internally validated for hyperparameter (alpha) tuning using a separate 10-fold cross-validation scheme within the training sample. Resulting models were tested by generating predicted scores for the held-out participants and performance was assessed by computing the partial correlation between predicted and observed scores. More specifically, partial correlation was used to account for head motion (mean frame-to-frame displacement; mean rotation; maximum displacement) and quantity of data (number of frames). Given the large sample size and likelihood of inflated parametric p-values, 10,000 bootstrapped samples were used to generate 95% confidence intervals around estimates to interpret model performance.

To consider the possibility that models generated from rest and task data complement each other in a way that cumulatively improves prediction, data-driven models were created using a combination of all tasks and rest. Vectorized functional connectivity matrices were concatenated for participants meeting inclusion criteria for all four scan conditions (resting-state, MID, SST, and n-back; n = 741) and subsequently used to predict component scores as previously described. Motion parameters (mean and maximum frame-to-frame displacement, maximum rotation) were averaged across all scan conditions to control for motion in the combined model. To account for the possibility that differences in model performance in the combined model could be due to a greater quantity of data used per subject, feature selection was performed in a separate model to select the number of features equal to a single task- or rest-based functional connectivity matrix. This feature selection step was performed by computing univariate correlations between edge strengths and component scores within the training set and selecting the top 35,778 edges for model construction.

#### Data visualization

The most informative connections for each model (top 0.1%) were arranged according to independently-defined functional networks (described in greater detail in (Finn et al., 2015) and visualized using MNE-Python’s *viz* package (Gramfort, 2013).

## Results

### Behavioral results

Pairwise correlations visualized across all substance use measures demonstrated a range of associations (Figure 2A). Multidimensional scaling of pairwise Euclidean distances amongst substance use data depicted a distinction between risk-seeking and familial factors (Figure 2B). Principal components analysis revealed two orthogonal dimensions that were highly reliable across 10,000 bootstrapped samples (Figure 2D). The first component (PC1) loaded primarily onto questions associated with early sips of alcohol (e.g., obtained without parental permission) as well as curiosity and intention to experiment with substances. The second component (PC2) loaded onto questions associated with family substance use, including a family history of substance use disorder and whether someone smokes in the child’s home. These two components accounted for 23% and 14% of the total variance, respectively (Figure 2C). Because of their high reliability, in addition to accounting for the most variance (37% cumulative), these first two components were of primary interest in subsequent analyses.

**Figure 2.**
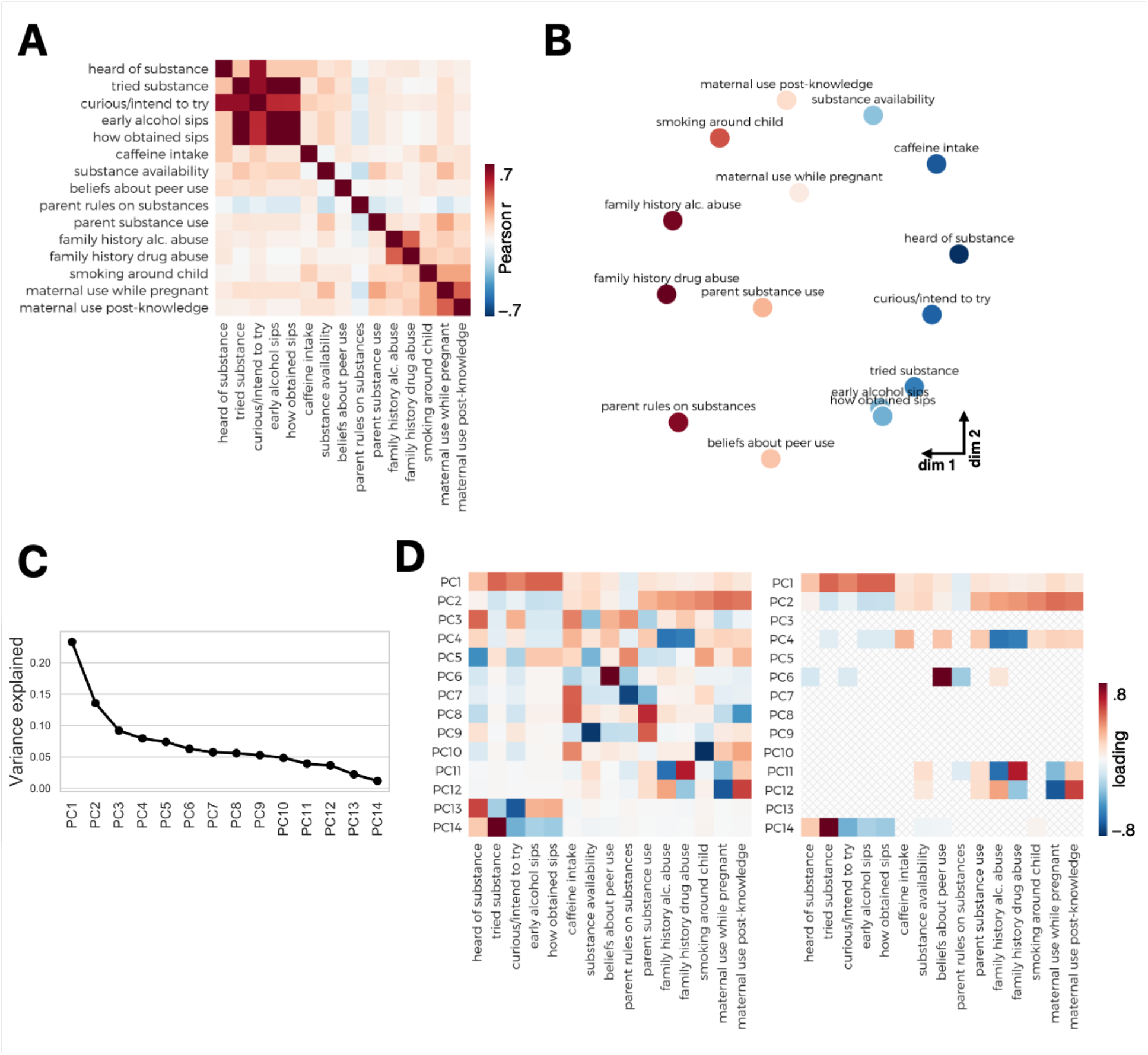
Substance use measures. **A)** Correlation matrix of transformed behavioral measures related to substances. **B)** MDS plot visualizing distances among substance use measures; **C)** Scree plot depicting variance explained by individual components. **D)** PCA loadings for each behavioral measure (left). PCA loadings that demonstrated consistency across 10,000 bootstrapped samples (right).

### Connectome-based prediction

Functional connectivity during tasks and rest was used to generate models predicting variability in PC1 (risk-seeking associated with substance us) and PC2 (familial risk factors). Individual differences in scores for the substance use risk-seeking component (PC1) were significantly predicted from connectivity during rest (partial correlation [r_p_] = 0.1, 95% bootstrap CI: [0.06, 0.13]) as well as during all three tasks (MID: r_p_ = 0.09, 95% CI [0.05, 0.14]; SST: r_p_ = 0.08, 95% CI [0.03, 0.13]; N-Back: r_p_ = 0.15, 95% CI [0.1, 0.2]) (Figure 3). Scores for familial risk factors associated with substance use were significantly predicted by functional connectivity during the emotional n-back task (r_p_ = 0.06, 95% bootstrap CI: [0.005, 0.11]), but not during rest, MID or SST. Findings were consistent when siblings were included in the analysis (Figure S3). Model performance was not associated with the number of participants included in a given scan condition (p = 0.86) suggesting that quantity of data did not bias performance of these models.

**Figure 3.**
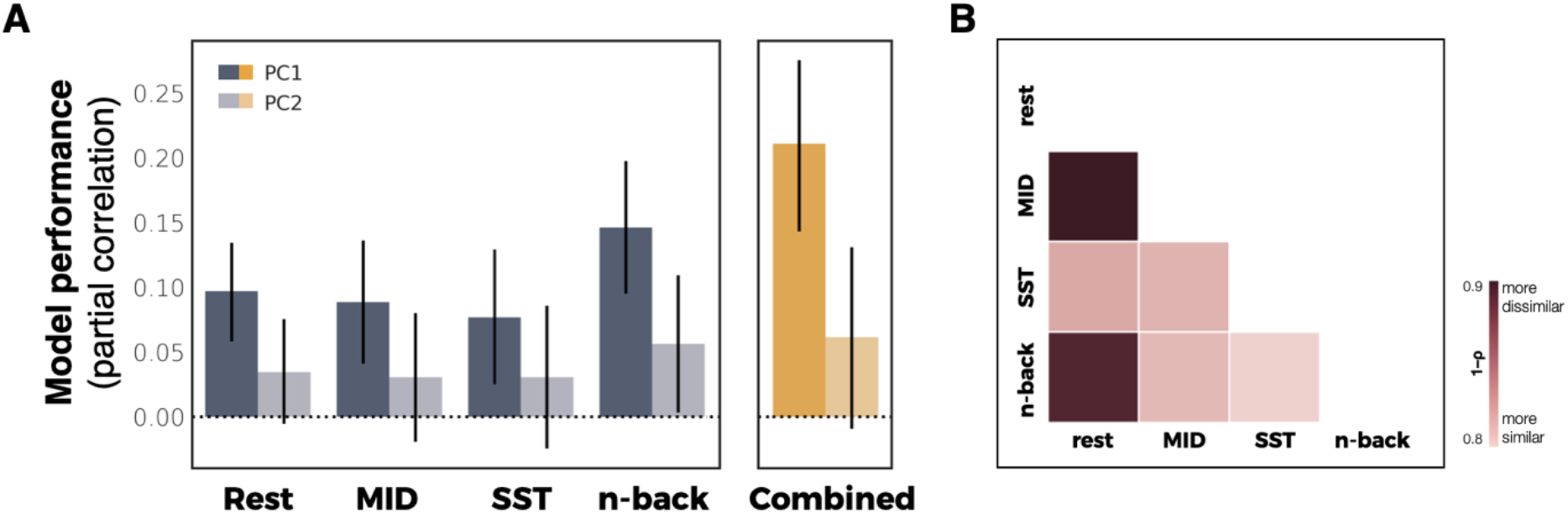
Predictive model performance. **A)** Cross validated predictions of substance-related risk components from functional connectivity. Model performance is defined as the partial correlation between predicted and observed values accounting for covariates. Error bars represent 95% confidence intervals. **B)** Pairwise distances (1 – Spearman correlation) between coefficients for all task- and rest-based models for PC1.

Combining all tasks improved model performance for predicting PC1 (r_p_ = 0.21, 95% bootstrap CI: [0.14, 0.28]) but not PC2 (r_p_ = 0.06, 95% bootstrap CI: [−0.01, 0.13]) (Figure 3). These results were consistent if the number of features selected matched that of a single scan condition (e.g., a single task, 35778 edges), with a marginal improvement in predicting PC2 (PC1: r_p_ = 0.19, 95% bootstrap CI: [0.12, 0.26]; PC2: r_p_ = 0.07, 95% bootstrap CI: [0.004, 0.14].

Mean frame-to-frame displacement across all tasks and rest was not correlated with PC1 (r = −0.01; p 0.45) or PC2 scores (r = 0.0005; p = 0.97), suggesting that head motion was not associated with behavioral phenotypes. Likewise, there was no relationship with maximum rotation and behavioral scores (PC1: r = 0.01, p = 0.5; PC2: r = −0.003, p = 0.85).

Although predictive models did not significantly differ in terms of performance, model coefficients demonstrated substantial differences across conditions and components. A quantitative measure of model distinctiveness (1 – Spearman correlation) demonstrated dissimilarity between rest and task conditions, with models derived from the MID task demonstrating the greatest dissimilarity with rest for both PC1 (1–ρ = 0.88; 95% CI: [0.87, 0.89]) (Figure 3B). Model coefficients were reliable across folds within task and rest conditions (mean [s.d.] correlation = 0.84 [0.014]). Visualizing the most informative connections (top 0.1%) of each predictive model further illustrates the distinctiveness across models (Figure 4).

**Figure 4.**
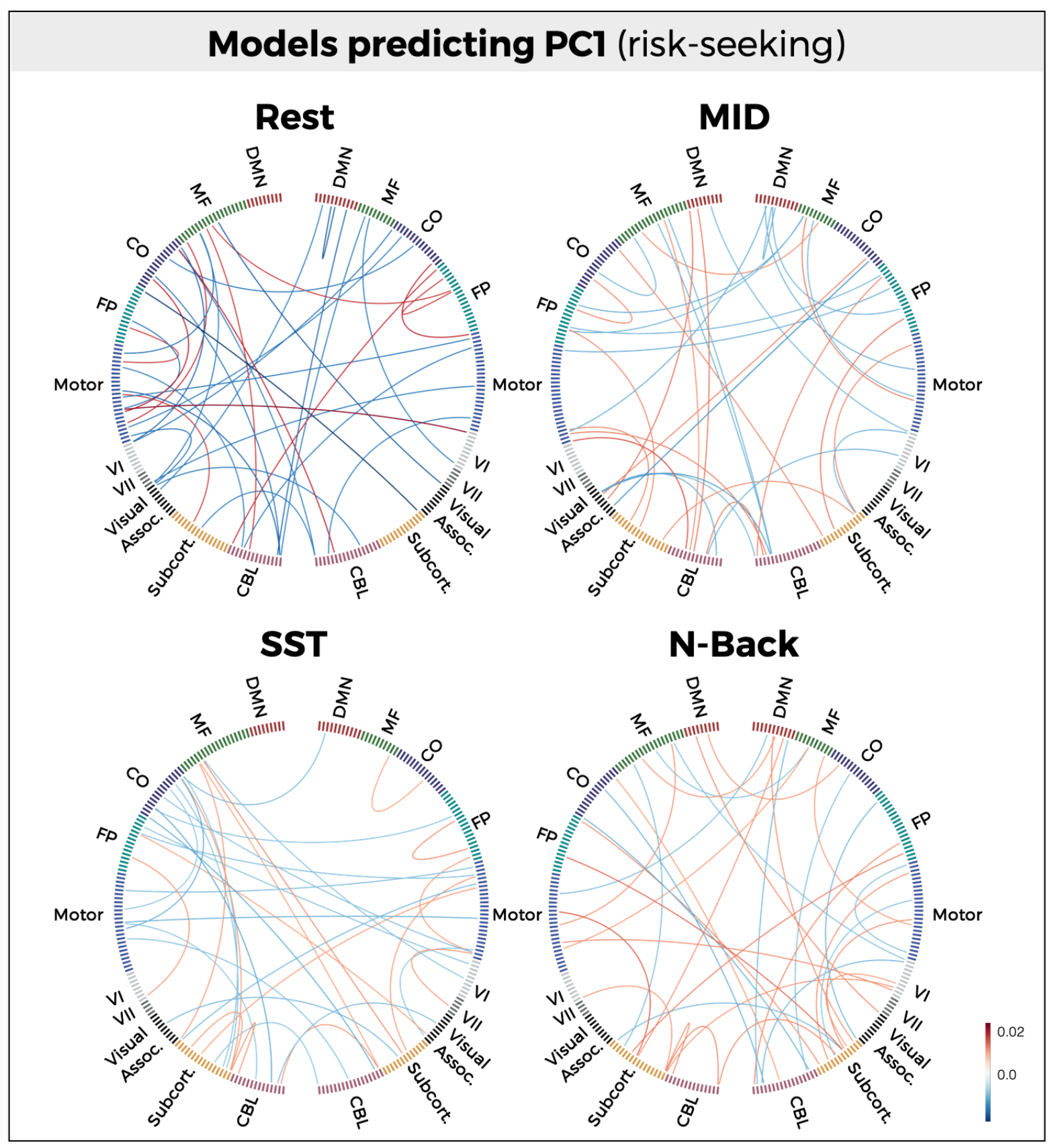
Anatomical specificity of predictive models. Most informative connections (top 0.1%) in connectome-based models predicting (A) PC1 (risk-seeking associated with substance use) and (B) PC2 (familial risk factors associated with substance use). Nodes are arranged in correspondence with networks functionally defined in an independent dataset. Edges represent model coefficient values. (DMN = Default mode network; MF = Medial frontal; CO = Cingulo-opercular network; FP = Frontoparietal network; CBL = Cerebellum)

## Discussion

Through the use of data-driven predictive models developed in a large cohort of children, the current study found an association between risk factors for substance use vulnerability and functional connectivity in the developing brain. Dimensionality reduction of a large battery of substance-related behavioral measures in 9,437 nine- and ten-year-olds revealed two latent dimensions of risk. The first loaded heavily onto risk-seeking (e.g., curiosity to try substances) and the second loaded on familial risk factors (e.g., family history of substance use). Regularized regression models trained on functional connectivity data significantly predicted these dimensions in a subset of participants (Year 1 Fast Track). Specifically, functional connectivity during a range of cognitive conditions, including rest and task, significantly predicted individual differences in PC1 (risk-seeking associated with substances). Predictive models were supported by highly distributed patterns of connections that varied across scan conditions. Consistent with evidence showing that a combination of task- and rest-based connectomes offer complementary information yielding greater prediction accuracy (Gao et al., 2019), predictive performance increased when all scan conditions were included in the model, independent of the number of features used. By contrast, functional connectivity was a weaker predictor of PC2 (familial risk factors). Connectivity during a general cognitive task (n-back) weakly—but significantly—predicted this latent dimension, whereas connectivity during tasks associated with reward motivation and cognitive control showed no predictive relationship with PC2.

These findings complement a growing body of literature aimed at identifying predictive measures of vulnerability to future substance use (Heitzeg et al., 2015; Squeglia & Cservenka, 2017; Tervo-Clemmens et al., 2020). One way in which brain-based predictions have been generated is through longitudinal and prospective approaches. For example, brain responses during an inhibitory control task have been found to differentiate adolescents that became heavy alcohol users from those that did not after 18 months (Mahmood et al., 2013), three years (Wetherill et al., 2013), and after four years (Heitzeg et al., 2014; Norman et al., 2011). Other prospective studies have related baseline brain responses to alcohol cues to frequency (Courtney et al., 2018) and severity (Dager et al., 2014) of alcohol misuse among college-aged students over the course of several months. Although these studies have provided insight into neural markers that predict future behavior, they may not capture variability that is specific to substance use vulnerability in at-risk populations. In comparison to prospective predictions of substance use behavior, other work has sought to identify biomarkers of vulnerability by examining associations with known risk factors for substance use. For example, previous work has observed differences in reward-related responses in drug-naive children with a family history of substance abuse (Ivanov et al., 2012) as well as in children with a family history of alcohol use disorder (Heitzeg et al., 2010; Yau et al., 2012).

The current study utilizes a unique and complementary approach to predicting substance use vulnerability. Whereas identifying common neural pathways or distinct behavioral risk factors will give rise to a greater understanding of neurobiological and psychological mechanisms underlying vulnerability, predictive models may afford greater generalizability (Yarkoni & Westfall, 2017). Further, mechanistic approaches require greater consideration of how to operationalize a given phenotype—such as vulnerability, which may lead to a more reductionist viewpoint that overlooks the broader complexity of that phenotype (Varoquaux & Poldrack, 2019). Predictive models have the potential to capture more highly distributed features contributing to a particular outcome (e.g., beyond canonical networks), and when constrained by hypothesis-driven approaches, may inform mechanistic perspectives of these features. Thus, both explanation-based and predictive-based approaches will be instrumental in establishing biomarkers of vulnerability, which may ultimately inform early detection of risk for addiction. Despite the utility of predictive modeling, the current study presents a first step towards uncovering the broader potential of this approach in understanding substance use vulnerability. As a result, there are a number of avenues for future work and limitations that warrant further discussion.

Although the current analysis restricted inclusion of behavioral variables to those directly related to substances, future work should address the role of co-occurring psychiatric and behavioral factors (e.g., impulsivity, sensation-seeking) associated with substance use. For example, a substantial body of evidence has linked externalizing psychopathology with early initiation and experimentation with substances (Colder et al., 2013; Iacono et al., 2008; King et al., 2004; White et al., 2001), with a 47% median prevalence of comorbidity across fifteen community samples of adolescents (Armstrong & Costello, 2002). Individuals with substance dependence and adolescents with externalizing disorders both demonstrate increased striatal activity during receipt of reward (Bjork et al., 2009; Luijten et al., 2017) as well as decreased cortical activity associated with inhibitory control (Luijten et al., 2014; Rubia et al., 2009; A. B. Smith et al., 2006). Moreover, reward-related activity has been shown to positively correlate with externalizing behaviors as well as drinking behavior in children genetically at risk for alcohol dependence (Yau et al., 2012), and similarly, control-related activity in children has been shown to prospectively predict externalizing behavior as well as later substance use (Heitzeg et al., 2014). These findings may imply a moderating role of externalizing psychopathology in substance use vulnerability (Bjork et al., 2017; Heitzeg et al., 2015). Consistent with this account, a recent meta-analysis found greater reliability of striatal activity associated with substance use vulnerability in adolescents when co-occurring externalizing disorders were present (Tervo-Clemmens et al., 2020).

Externalizing behaviors and risk for addiction (Krueger et al., 2007) have been suggested to have a common neurogenetic liability, wherein heritable features associated with behavioral disinhibition are expressed in brain circuitry motivating substance use (Iacono et al., 2008). Studies performed in adolescent twin pairs support this theory by demonstrating moderate heritability of externalizing disorders (Cosgrove et al., 2011) as well as a highly heritable dimension of behavioral disinhibition that explained approximately half of the variability in substance use (Young et al., 2000). Together these findings suggest that vulnerability to substance use and externalizing psychopathology are heritable and are represented by a common neurogenetic liability underlying comorbidity.

Consistent with this work, the latent behavioral dimensions of vulnerability identified in the current study were significantly correlated with externalizing symptoms. In particular, the familial factors dimension (PC2) explained approximately 5% of the variance in externalizing scores (r = 0.22), whereas the risk-seeking dimension (PC1) explained only 0.6% (r = 0.08) (Figure S4). The stronger relationship between externalizing and familial risk factors is consistent with literature describing a shared heritable component between externalizing psychopathology and substance use vulnerability. By contrast, the weak relationship between externalizing scores and the risk-seeking dimension suggests that variability associated with externalizing psychopathology is not heavily contributing to or biasing the predictive models derived from PC1. Future work should further explore the relationship between externalizing and other behavioral risk factors with neurobiological vulnerability to substance use.

The current study provides insight into the potential for generating data-driven models that prospectively predict behavioral outcomes prior to knowledge about those outcomes. An open question is whether the models of vulnerability generated here differentially predict early initiation and future substance use during adolescence and young adulthood, and further, how these models may capture differences in behavior across development. Twin studies have demonstrated a dynamic pattern of behavioral disinhibition across development, such that the heritability of substance use decreased from 58% at age 12 to 20% by age 17 (Young et al., 2009). This finding suggests the possibility that heritable factors play a greater role in the initiation of early substance use, whereas environmental factors may play a greater role in continuation and severity of use. Indeed, other studies in twins report early alcohol consumption to be a risk factor for later dependence, independent of genetic or shared family environmental factors (Grant et al., 2006). Thus, one possibility may be that the second risk dimension found here (PC2; familial factors) will predict experimentation or early initiation of substance use, and that the first dimension (PC1; risk-seeking) and corresponding connectome-based models will predict continuation or severity of use. As the ABCD Study continues to collect longitudinal data, these questions and hypotheses—including disentangling the differential contributions of psychopathology, substance use, and heritability across development (Bjork et al., 2017)—can be further tested and explored.

Another interesting avenue for future research concerns the extent to which different scan conditions may predict later outcomes. The current study found no meaningful differences among task conditions and rest, such that all conditions similarly predicted the risk-seeking dimension (PC1) and failed to predict (or weakly predicted) familial factors associated with substance use (PC2). Given prior work identifying correlates between substance use and different brain regions involved in reward, cognitive control, and working memory, it will be important to test the possibility that certain tasks or rest conditions may better predict future behavior. With that said, comparisons of task conditions and rest will require additional consideration of covariates that may contribute technical differences across scan conditions. For example, the reliability of individual measures of functional connectivity increases as a function of data quantity (Gordon et al., 2017). Another consideration will be to examine how potential task-driven effects differ from the underlying connectivity and to what extent these effects bias model performance. The current analysis did not remove task-induced fluctuations from the task-based data given work demonstrating that these fluctuations may enhance individual differences and improve prediction (Finn et al., 2017; Greene et al., 2018; Rosenberg et al., 2016a; Yoo et al., 2018); however, it will be interesting to examine how potential task-driven effects differ from the underlying connectivity and to what extent these effects bias model performance in future work.

The current study is well-powered to detect the effects observed here; however, the modest size of these effects is an important limitation to consider. The bias to publish papers with large effect sizes in small sample sizes has been well-documented (Ioannidis, 1998; Sterne et al., 2000), including one study that observed a highly significant negative correlation between sample size and effect size in more than 1,000 psychological studies (Kühberger et al., 2014). In concert with evidence demonstrating that larger sample sizes tend to yield smaller effect sizes (S. M. Smith & Nichols, 2018)—including within the ABCD Study (Marek et al., 2020)—and conversely, that statistically significant effects in underpowered studies are likely to be inflated (Button et al., 2013; Ioannidis, 2008; Marek et al., 2020; Yarkoni, 2009), it is unsurprising that the effect sizes observed in the current study are small (partial correlation [r_p_] ~= 0.1). However, a small effect in a large sample size potentially allows for higher replicability (Marek et al., 2020; Scheinost et al., 2019), which is further demonstrated here via cross-validation procedures and testing on novel participants. Finally, it is worth noting that the effects reported here are observed within children that have yet to begin using substances, and therefore effect sizes of this scale are not unexpected. In addition to leveraging models at baseline to generate prospective predictions, future work should aim to address developmental changes in the performance of predictive models (i.e., effect sizes) in order to assess the strength and emergence of these associations over time.

Another potential limitation of the current study concerns specific methodological considerations. For example, the current analysis used PCA to reduce the dimensionality of the behavioral data, which required transformation to reduce the skewness of the data. Future work will be needed to assess the robustness of this data reduction step relative to other techniques such as polychoric correlation, which may be better suited for variables of mixed types (i.e., ordinal, numerical) (Holgado–Tello & Chacón–Moscoso, 2010). Moreover, the number of participants lost to motion were substantial—in part due to rigorous motion thresholds often applied to adult data (Horien et al., 2018; Kumar et al., 2019; Rosenberg et al., 2016a; Rosenberg et al., 2016b), which is similarly observed by others using rigorous quality control procedures within ABCD (Marek et al., 2020; Sripada et al., 2019) and in connectivity-based studies in adults and adolescents (Greene et al., 2018; Horien et al., 2019). Analytic tools such as censoring (Power et al., 2014) may prevent data loss due to motion, which may subsequently allow for greater variability in the distribution of behavioral variables of potential interest that may be related to motion (e.g., impulsivity). Finally, greater consideration should be given to the role of demographic covariates in the current models. The current analysis controlled for motion and amount of data; however, age, sex, race, and data collection site were not included in these models. Although this analysis may suggest that the models identified here are generalizable across demographic factors (i.e., through random sampling across folds), future work is needed to explicitly test to what extent these variables impact model performance.

In contrast to studies that have focused on specific risk factors and/or risk for specific substances, the current study followed a data-driven approach that allowed for the identification of latent dimensions of substance-general vulnerability. In doing so, we captured and predicted two dimensions that represent distinct patterns of information in both the brain and behavior. These findings extend prior work on the role of developing reward- and control-related circuitry in motivating adolescent risk seeking and substance use initiation, and provide insight into the potential heritability of familial risk factors for substance use. Ultimately, predictive models of substance use vulnerability may inform early identification and addiction prevention strategies in at-risk children.

## Acknowledgements

This work was funded by NIH U01 DA041174 (awarded to B.J.C.).

