## Supplemental materials for "Behavioral and brain signatures of substance use vulnerability in childhood"

| Assessment | NDA variables | Description | Example questions used |
|---|---|---|---|
| **ABCD Youth Substance Use Interview** Lifetime use (Lisdahl & Price, 2012)**;** TLFB (Sobell et al., 1996); iSip (K. M. Jackson et al., 2015); Peer deviance (Lloyd D. Johnston et al., 2017) | ***abcd_ysu02;*** *tlfb_*;* *path_*;* **isip_*;* *peer_deviance** | Curiosity and intent to use substances; low-level alcohol, cannabis, and nicotine use; acute subjective response to alcohol, tobacco, and marijuana; perceptions of substance use in peer group | **Timeline follow back (TLFB)**<br>*"Have you [heard of / tried] __";* // 0 = No; 1 = Yes<br><br>**Pop. assessment of tobacco & health (PATH)**<br>*"Are you curious to try __?"* // 1 = Very curious; 2 = Somewhat curious; 3 = A little curious; 4 = Not at all curious; 5 = Don't Know; 6 = Refused<br>*"Do you think you will try __ soon?"* // 1 = Definitely yes; 2 = Probably yes; 3 = Probably not; 4 = Definitely not; 5 = Don't know; 6 = Refused<br><br>**iSip**<br>*"Have you ever had alcohol (not as part of a religious ceremony)?, "How did you obtain it?"* // 0 = No; 1 = Yes \| 1 = My mom; 2 = My dad; 3 = Other guardian; 4 = An uncle or aunt; \| 1 = They offered me a sip; 2 = I intentionally took it when they weren't looking; 3 = I accidentally took it when they weren't looking<br><br>**Peer deviance**<br>*"How many of your friends use __?"* // 0 = None; 1 = A Few; 2 = Some; 3 = Most; 4 = All |
| **ABCD Summary Scores Substance Use** | ***abcd_suss01;*** *su_caff_ss_sum_calc* | Caffeine summary scores | *"How many [caffeinated] drinks did you have per week in the past month?"* // Enter numerical value |
| **ABCD Parental Rules on Substance Use** (Arthur et al., 2007) | ***prq01;*** *parent_rules_q** | Parental substance use approval and rules (alcohol, cigarette, and marijuana use) | *"Which statement best describes the rules for your child's drinking in your house?"* // 1 = My child is not allowed to drink under any circumstances; 2 = My child is allowed to drink at home but only on special occasions and under parent supervision; 3 = My child can have a drink when he/she asks for one; 4 = My child can drink at home whenever he/she wants to; 5 = I don't set rules about my child's drinking; 6 = I haven't made rules yet about my child drinking<br><br>*"Are these rules the same for all family members?"* // 1 = No, applies only to children, not teen or adults; 2 = No, applies only to children and teen, not adults; 3 = Yes<br><br>*"Are there penalties for violating family rules about drinking?"* // 0 = No; 1 = Yes |
| **ABCD Parent Community Risk and Protective Factors** (Lloyd D. Johnston et al., 2017) | ***abcd_crpf01;*** *su_risk_p_** | Beliefs about drug accessibility and availability (alcohol, nicotine, marijuana, and 'other' drugs) | *"If your child wanted to get some [substance] how easy would it be for her/him to get some?"* // 0 = Very hard; 1 = Sort of hard; 2 = Sort of easy; 3 = Very easy; 4 = Don't know |

| | | | |
|---|---|---|---|
| **ABCD Family History Assessment Part 1** | ***fhxp102;*** *fam_history_5*;* *famhx_4_p* | Family history of psychopathology and substance use | *"Has ANY blood relative of your child ever had any problems due to alcohol?"* // 0 = No; 1 = Yes<br><br>*"Has ANY blood relative of your child ever had any problems due to drugs?"* // 0 = No; 1 = Yes |
| **ABCD screener** | ***abcd_screen01;*** *scrn_hr_smoke* | Eligibility screener and screener risk measures | *"In the past six months, has anyone who lives with your child smoked a cigarette, e-cigarette, or used any other type of tobacco?"* // 0 = No; 1 = Yes |
| **ABCD Parent Adult Self Report Raw Scores Aseba (ASR)** | ***pasr01;*** *asr_q06_p;* *asr_q90_p;* *asr_q124_p;* *asr_q126_p* | Parent dimensional psychopathology | *"I use drugs (other than alcohol, nicotine) for nonmedical purposes."* // 0 = Not True; 1 = Somewhat/Sometimes True; 2 = Very True/Often True<br><br>*"I drink too much alcohol or get drunk."* // 0 = Not True; 1 = Somewhat/Sometimes True; 2 = Very True/Often True<br><br>*"In the past 6 months, about how many times per day did you use tobacco."* // Numerical value (0::100; integer)<br><br>*"In the past 6 months, on how many days did you use drugs for nonmedical purposes."* // Numerical value (0::182; integer) |
| **ABCD Developmental History Questionnaire** | ***dhx01;*** *devhx_8_*;* *devhx_9_** | Prenatal exposure before and during pregnancy (medications, drugs alcohol, tobacco) | *"Before knowing of pregnancy. Alcohol? Maximum drinks in one sitting? Average drinks per week? How many drinks did it take to feel the effects?"* // 1 = Yes; 0 = No<br><br>*"After knowing of pregnancy. Alcohol? Maximum drinks in one sitting? Average drinks per week? How many drinks did it take to feel the effects?"* // 1 = Yes; 0 = No |

**Table S1. Descriptions of substance use-related questionnaires and behavioral measures.** NIMH Data Archive (NDA) names list variables used for each assessment.

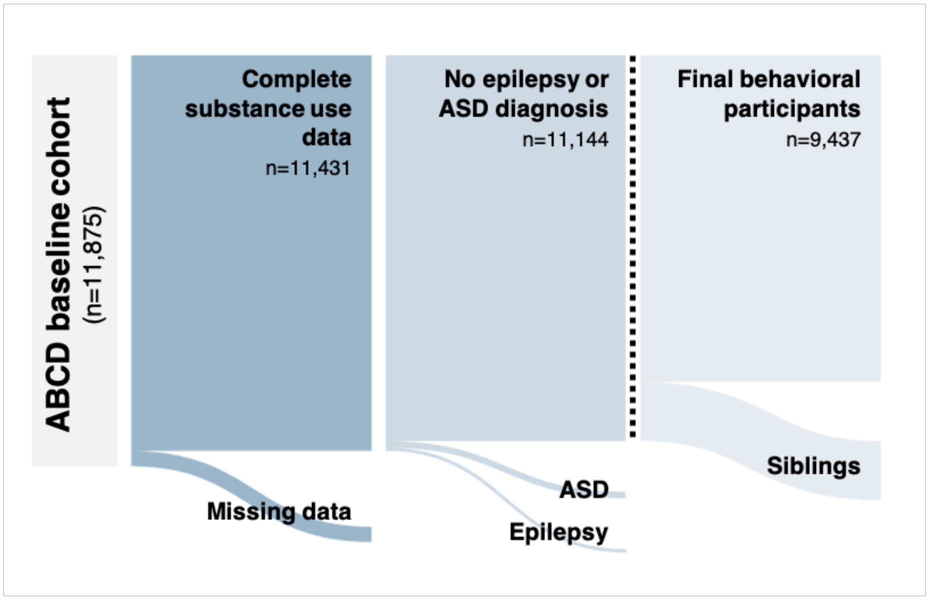


**Figure S1. Behavioral participant selection.** Sankey diagrams illustrating selection procedure for behavioral analyses, including exclusion criteria based on data completeness and participant demographics. Dotted line indicates pre- and post-exclusion of family relatedness used in analyses with and without siblings.

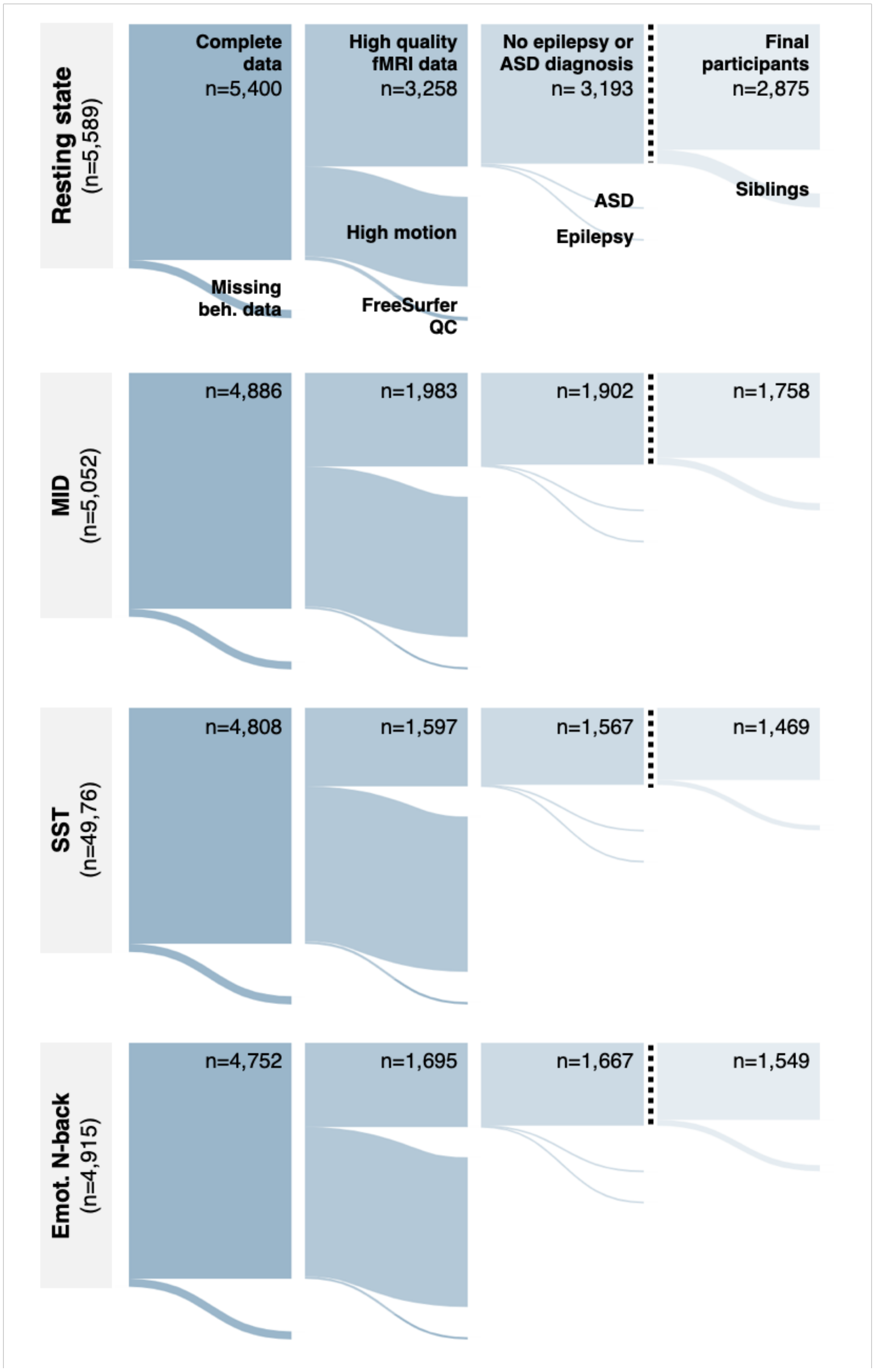


**Figure S2. Neuroimaging participant selection.** Sankey diagrams illustrating selection procedure for each scan condition, including exclusion criteria based on data completeness, data quality, and participant demographics. Dotted line indicates pre- and post-exclusion of family relatedness used in analyses with and without siblings. (QC = quality control)

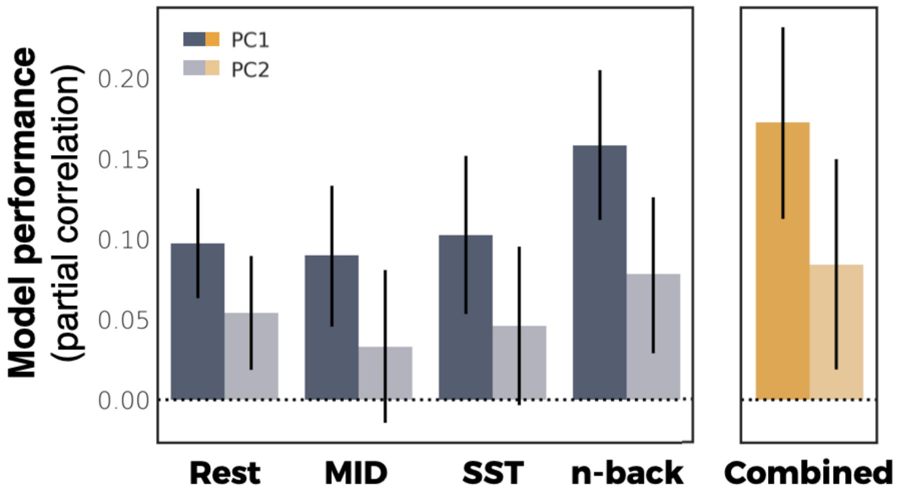


**Figure S3. Predictive model performance when including siblings.** Cross validated predictions of substance-related risk components from functional connectivity. Model performance is defined as the partial correlation between predicted and observed values accounting for covariates. Results are consistent with models excluding siblings with the exception of rest (r = 0.05, 95% CI = [0.02, 0.09]) and the "combined" models (r = 0.08, 95% CI = [0.02, 0.15]) predicting PC2. This improvement is likely due to the presence of siblings impacting prediction of familial factors associated with substance use.

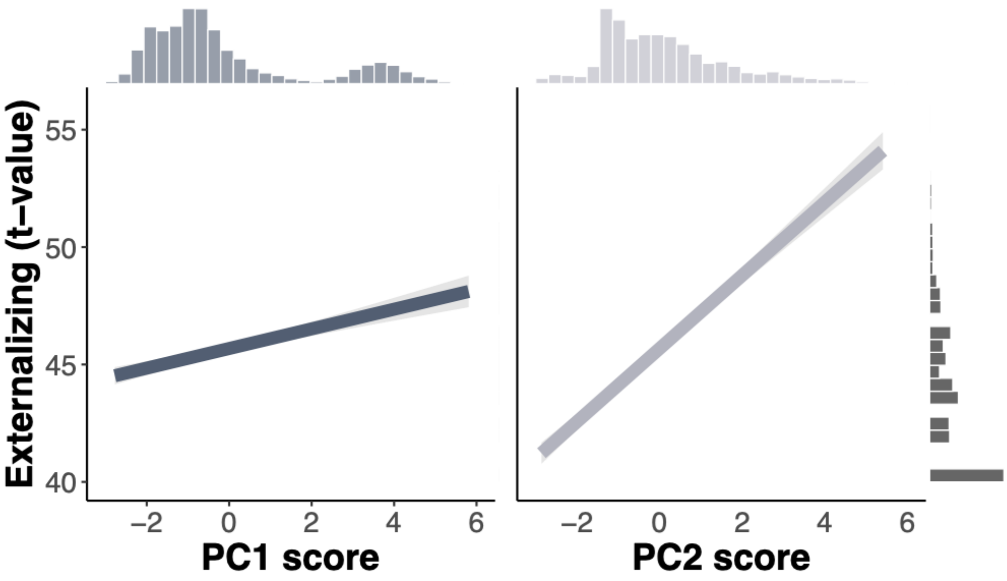


**Figure S4. Behavioral associations with externalizing symptoms.** Normed externalizing symptoms are correlated with PC1 (left; $r = 0.08$; non-parametric $p < 0.001$) and PC2 (right; $r = 0.22$; non-parametric $p < 0.001$) scores. Marginal histograms demonstrate distributions of scores on corresponding axes. Externalizing scores are adjusted for age and gender (NDA variable name: *cbcl_scr_syn_external_t*).

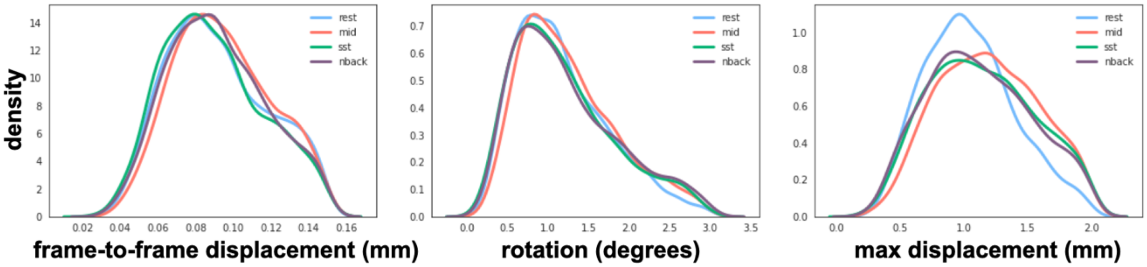


**Figure S5. Distribution of motion parameters.**

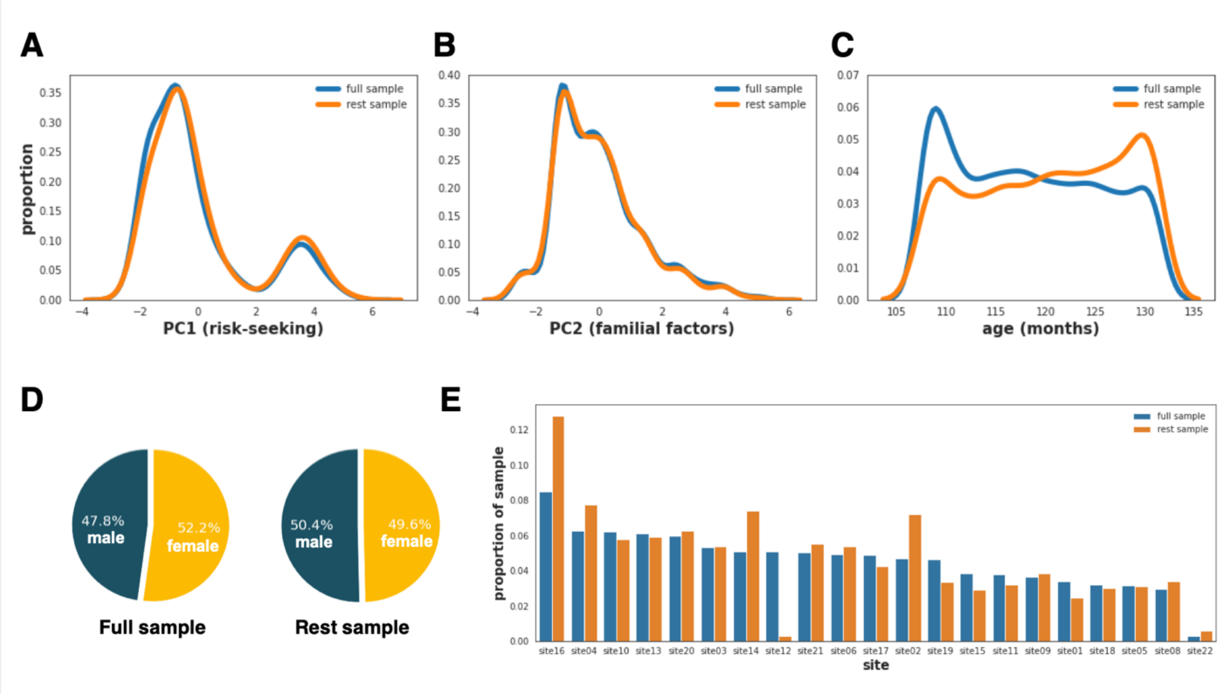


**Figure S6. Sample distributions comparing resting-state fMRI sample and the full sample (excluding resting-state participants).**
